# Creating something out of nothing: Symbolic and non-symbolic representations of numerical zero in the human brain

**DOI:** 10.1101/2024.01.30.577906

**Authors:** Benjy Barnett, Stephen M. Fleming

## Abstract

Representing the quantity zero as a symbolic concept is considered a unique achievement of abstract human thought. Despite considerable progress in understanding the neural code supporting natural numbers, how numerical zero is encoded in the human brain remains unknown. We find that both non-symbolic empty sets (the absence of dots on a screen) and symbolic zero (‘0’) occupy ordinal positions along graded neural number lines within posterior association cortex. Neural representations of zero are partly independent of numerical format, exhibiting distance effects with countable numerosities in the opposing (symbolic or non-symbolic) format. Our results show that format-invariant neural magnitude codes extend to judgements of numerical zero and help motivate theoretical accounts in which representations of symbolic zero are grounded in more basic representations of sensory absences.

## Introduction

The number zero plays a central role in science, mathematics, and human culture^1,2^ and its symbolic representation is considered a unique property of abstract human thought^2,3^. The psychological basis of zero is unusual: while natural numbers correspond to the observable number of countable items within a set (e.g., one bird; three clouds), an empty set does not contain any countable elements. To conceptualise zero, one must instead abstract away from the (absence of) sensory evidence to construct a representation of numerical absence: creating ‘something’ out of ‘nothing’^2,4,5^. Given these differences, it remains an open question as to how zero is represented in relation to other numbers.

In contrast to zero, the neural representation of natural numbers is better understood. Distinct neural populations are selective for specific numerosities, exhibiting overlapping tuning curves with neighbouring populations tuned to adjacent numerosities^6,7^. This architecture underpins a so-called distance effect^8^, where numbers close together in numerical space have similar neural representations. For instance, neural responses to numbers one and two are more similar than neural responses to one and ten^7,9,10^. Importantly, a component of this neural code is thought to be invariant to numerical format^11–15^ such that, for example, neural representations of ‘six’ are shared across symbolic and non-symbolic formats (e.g., both the Arabic numeral ‘6’ and six dots; although see^16^). In humans, these format-invariant representations of numerical magnitude have been localised to the parietal cortex^11,13,14^, with topographic maps underpinning numerosity perception found more broadly across association cortex^17,18^.

Although behavioural evidence suggests that zero occupies a place at the beginning of this mental number line^19–21^, zero is also associated with unique behavioural and developmental profiles compared to natural numbers. For instance, the reading times of human adults are increased for zero compared to non-zero numbers^39^ and zero concepts emerge later in children than those for natural numbers^5,46,47^. Distinct behavioural characteristics associated with zero are not surprising given the heightened degree of abstraction required to conceptualise numerical absence. In turn, it is plausible that neural representations of zero are distinct to the scheme that has been discovered for natural numbers^25^. Initial research in non-human animals has indicated that numerical zero shares some neural properties with natural numerosities, such as overlapping tuning curves and associated distance effects, along with invariance to particular stimulus properties^22–24^. Moreover, behavioural evidence for zero representations has been reported in a number of animals, including macaque monkeys^30^, African grey parrots^32^, and honeybees^34^. However, it remains unknown whether the symbolic, human conceptualisation of numerical zero, which emerges later than natural numbers in human culture^1,26^, engenders representations of zero that are both distinct from other numbers and which studies in non-human animals may have failed to reveal.

We tackled this question by employing two qualitatively different numerical tasks in humans while leveraging methodological advances to reveal the representational content of neural responses to numerical stimuli in magnetoencephalography (MEG) data^10,37^. Our choice of tasks was guided by previous work examining both non-symbolic^24^ and symbolic^10^ numerosity representations. Importantly, the use of two distinct tasks required participants to adopt distinct mathematical attitudes towards zero, ensuring that any commonalities between symbolic and non-symbolic neural representations of zero were not confounded by task-related processing. We assay both neural representations of non-symbolic numerosities (dot patterns), including zero (empty sets), and symbolic numerals, including symbolic zero. Numerosities ranged from zero to five, allowing us to examine the fine-grained representations of numbers close to zero. Our results reveal that neural representations of zero are situated along a graded neural number line shared with other natural numbers. Notably, symbolic representations of zero generalised to predict non-symbolic empty sets. We go on to localise abstract representations of numerical zero to posterior association cortex, extending the purview of parietal cortex in human numerical cognition to encompass representations of zero^14,18^.

## Results

Twenty-nine human participants (24 after exclusions; see Methods for details) took part in a magnetoencephalography (MEG) experiment involving two numerical tasks. The first was a non-symbolic match-to-sample task (Figure 1A) where participants observed two sequentially presented dot patterns that ranged in number from zero dots (empty set) to five dots^24^. Participants were asked to report whether the patterns contained the same or different number of dots. We employed two sets of dot patterns: a standard set which randomised the size of dots within each pattern, and a control set which kept total dot area, density, and luminance constant across numerosities (Figure 1D). To ensure participants could not rely on low level visual cues in identifying empty set stimuli, the background luminance was varied within and across stimulus sets, the background square size was randomly varied across all stimuli, and 50% of dots were of opposite contrast (white rather than black). The second task was a symbolic averaging task^10^ (Figure 1B). Here, participants observed a rapid serial presentation of 10 symbolic numerals from zero to five (0, 1, 2, 3, 4, 5), divided into orange and blue sets (5 numbers in each). Participants were asked to report the set of numbers with the higher or lower average. Decision type (higher or lower) was counterbalanced across participants. Employing different tasks per each notational format with different task requirements and decision types also ensured neural patterns induced by the perception of zero are unlikely to be driven by specific task features or calculation requirements. All analyses were exploratory and were not pre-registered prior to data collection.

**Figure 1.**
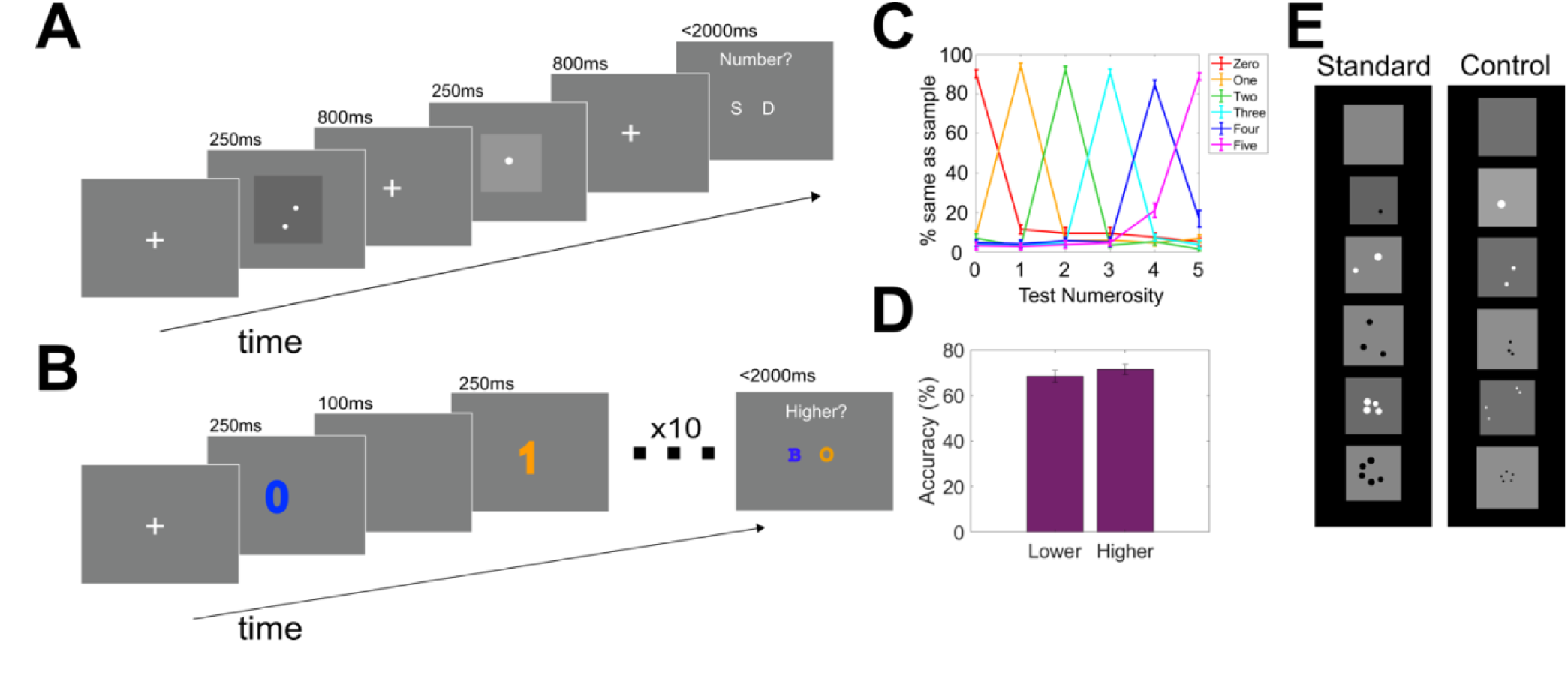
Experimental Procedure. *A. Trial structure for the non-symbolic match to sample task.* Participants observed a sample dot pattern followed by a test dot pattern before reporting whether the two patterns had the same or different numbers of dots. *B. Trial structure for the symbolic averaging task.* Participants observed a sequence of blue and orange numerals before reporting which set of numerals had the higher or lower average. *C. Behavioural tuning curves in the non-symbolic task.* Each curve reflects the percentage of trials that participants judged the test numerosity to be the same as the sample numerosity. Each colour represents trials with specific sample numerosities. The peak of each curve illustrates correct performance when the sample and test numerosities matched. Data points either side of the peak represent non-match trials. Error bars indicate SEM. *D.* Accuracy in the symbolic task split across participants who judged which set of numbers was higher, and those who judged which was lower. *E. Stimulus sets for non-symbolic task*. Dot size was pseudorandomised in the standard set, while low level properties of the dots including total dot area, density, and luminance were held constant in the control set. Across both sets, frame size of the dot patterns was randomly varied, to limit reliance on visual cues when identifying empty sets.

In the non-symbolic match-to-sample task, participants accurately determined whether dot patterns had the same or different numbers of dots (*Mean_accuracy_*: 0.92, *SE*: 0.16). Plotting behavioural tuning curves revealed near-ceiling performance across all numerosities (Figure 1C), with the exception of five-dot patterns which were more often confused with four-dot patterns than three-dot patterns (t(23) = 4.97, *p* < .001) – consistent with numerosity tuning curves becoming wider as number increases^8^. In the symbolic task, participants could reliably perform the task regardless of whether they were reporting the higher (*Mean_accuracy_*: 0.71, *SE*: 0.23) or lower (*Mean_accuracy_*: 0.68, *SE*: 0.27) average (Figure 1D), and there was no difference between performance across decision types (t(22) = -0.88, *p* = 0.39). As expected, performance was significantly higher in the non-symbolic match-to-sample task compared to the symbolic averaging task (t(23) = 8.82, *p* < .001).

### Identifying Neural Representations of Number

We next asked whether neural patterns recorded by MEG were sensitive to numerosity, by timelocking our data to the presentation of the dot pattern/symbolic numeral stimuli. Multiclass decoders were trained to classify different numerosities (zero to five) in both the non-symbolic and symbolic tasks. The frequency with which the decoders confused numerosities for one another is illustrated in Figure 2A. Here, individual panels represent trials where a particular numerosity was presented to the classifier, and the coloured lines indicate the proportion of those trials where the classifier predicted each one of the possible classes (zero to five) over the trial epoch. For example, the ‘NS-one’ panel shows that when one dot is presented in the non-symbolic task, the classifier predominantly and correctly labels this stimulus as numerosity one (yellow curve), with the next most likely error being a misclassification as the number two (green curve). Across all numbers and both formats, the classifiers successfully predicted the numerosity participants were viewing from their neural data, including zero numerosities (time-points where classifier significantly exceed chance level: non-symbolic: 70ms – 800ms; symbolic: 56.7ms – 800ms).

**Figure 2.**
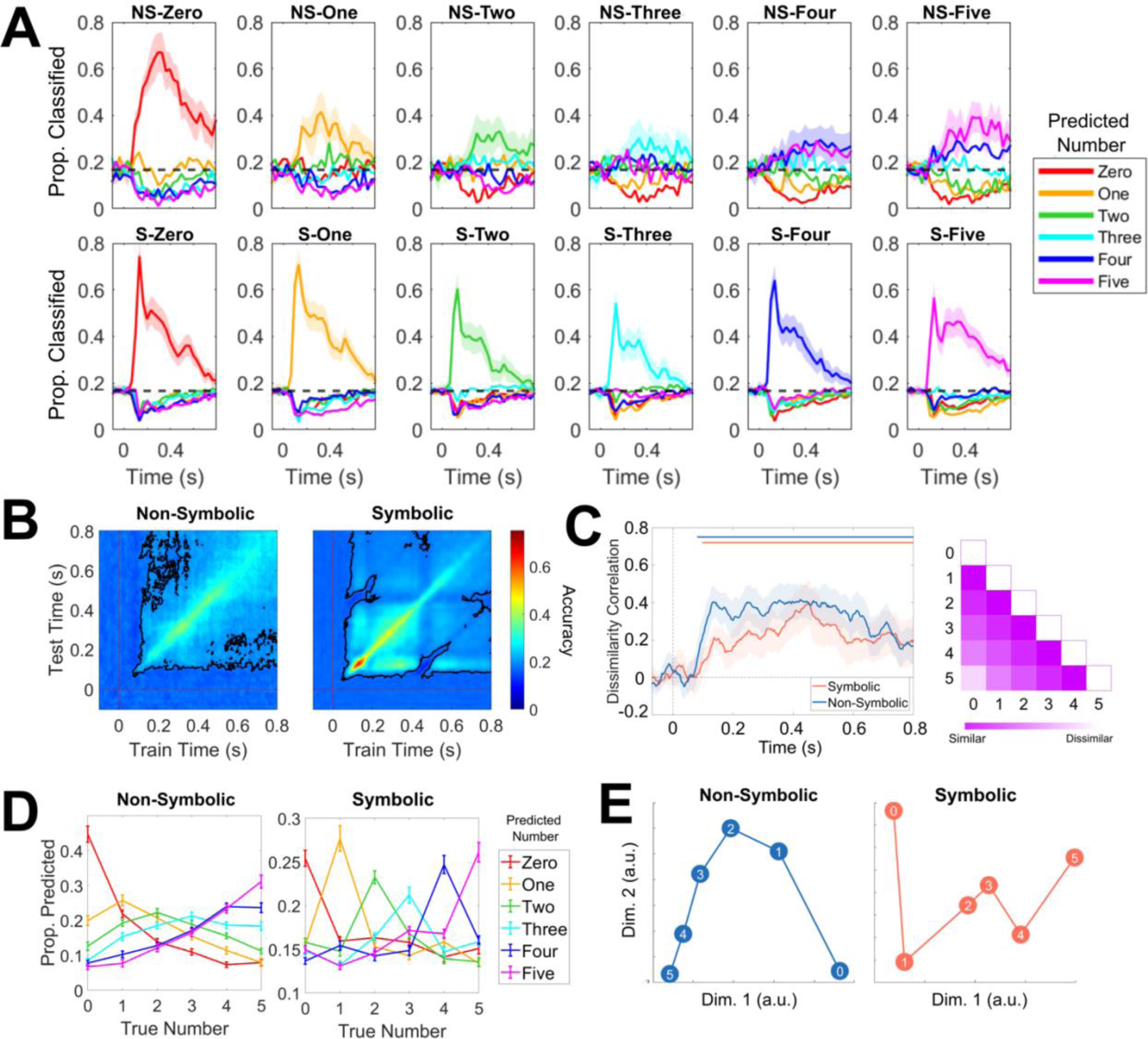
A Neural Number Line from Zero to Five. *A*. Across-time confusion matrices for multiclass decoders classifying non-symbolic (top) and symbolic numerosities (bottom). Individual panels represent trials where particular numerosities were presented to the classifier. Coloured lines indicate the proportion of those trials where the classifier predicted each numerosity. *B*. Temporal generalisation of multiclass decoders trained to decode numerosities zero to five in the non-symbolic (left) and symbolic (right) task reveals stable numerical representations over time in both tasks emerging shortly after stimulus presentation. Black lines illustrate timepoints where decoding was significantly above chance (*p* <.05, corrected for multiple comparisons). These stable time windows were used in the time-averaged analyses depicted in panels D and E. *C:* A model representational dissimilarity matrix (RDM) describing a distance effect from zero to five significantly predicted neural data in both non-symbolic and symbolic tasks. The diagonal of the RDM was not included in this analysis, preventing the self-similarity of each number from trivially explaining our results. Shaded areas indicate 95% confidence intervals. Horizontal lines show clusters of time where dissimilarity correlations were significantly above 0, *p* <.05 corrected for multiple comparisons. *D. Population level tuning curves derived from decoder confusion matrices.* Each curve represents the proportion of trials the classifier predicted a particular numerosity (indicated by the curve’s colour) as a function of the numerosity the decoder actually saw. For example, the red curve illustrates how the prediction of numerosity zero is distributed across different presented numerosities. For non-symbolic numerosities, the classifier confused numbers as a function of their numerical distance, consistent with a graded representation of numerical magnitude. In the symbolic task, representations were more categorical than the non-symbolic task. Error bars represent SEM. *E.* Multidimensional scaling of numerical representations in both tasks revealed a principal dimension which tracks numerical magnitude and a second dimension distinguishing extreme values from intermediate values.

We next leveraged temporal generalisation analysis to ask whether numerosity representations were stable over time^27^. When training and testing on all combinations of time points, stable time-windows where numerical information could be decoded above chance level were identified in both tasks from shortly after stimulus presentation up until the end of the analysed time window. This analysis was also used to generate Figure 2A, such that time-points at which classifiers exceed chance level are identical (non-symbolic: 70ms – 800ms; symbolic: 56.7ms – 800ms). These time windows in which stable numerosity representations were identified were used to create time-averaged data for use in subsequent population tuning curve (Figure 2D) and multidimensional scaling (Figure 2E) analyses.

### A Neural Number Line from Zero to Five

A fundamental feature of neural codes for natural numbers is a distance effect, whereby numbers closer together in numerical space are closer together in representational space^8,28^. Here we asked whether numerical zero exhibits similar distance effects with other numbers, consistent with it sharing a neural number line with countable numerosities. A Representational Similarity Matrix (RDM) describing a distance effect from zero to five successfully predicted neural data across both non-symbolic and symbolic numerical formats (Figure 2A). In the non-symbolic task, an RDM generalising numerical information across the two non-symbolic stimulus sets significantly predicted neural responses throughout the trial, indicating that neural correlates of number were independent of the physical properties of the dot stimuli (Supplemental Figure 2).

Multidimensional scaling of neural representations of numerosity in turn illustrates a distance effect (Figure 2E), with the numbers zero to five occupying positions along a single, ordered dimension, while a second dimension loosely distinguished intermediate numerosities (one to four) from the extremes (zero and five).

A stronger test of a distance effect in neural data is furnished by examining the confusability between neighbouring numerosities using population tuning curves (Figure 2D). These plots are time-averaged versions of the classifier confusion matrices in Figure 2A, i.e., the proportion of trials where the classifier predicted a particular numerosity as a function of the true numerosity within the time window in which numerical information could be reliably decoded (Figure 2B). For example, the red curve in Figure 2D indicates that the proportion of trials predicted as being zero peaks when the numerosity seen by the decoder was also zero, is next highest when the numerosity seen by the decoder was one, and so on.

In the non-symbolic task (Figure 2D, left), the classifier confuses zero with one (*Mean_propotion predicted_* = 0.218) more often than it confuses zero with two (*Mean_propotion predicted_* = 0.138) (t(23) = 6.23, *p* < .001). Similarly, it confuses one with two (*Mean_propotion predicted_* = 0.206) more often than with three (*Mean_propotion predicted_* = 0.155) (t(23) = 4.76, *p* < .001). This pattern of results is indicative of a gradedness in the representation of numerical magnitude across non-symbolic numerosities. In contrast, in the symbolic task (Figure 2D, right), the multiclass classifier does not confuse zero with one (*Mean_propotion predicted_* = 0.159) significantly more than it confuses zero with two (*Mean_propotion predicted_* = 0.163) (t(23) = -0.61, *p* = 0 .54), nor does it confuse one with two (*Mean_propotion predicted_* = 0.153) significantly more than it confuses one with three (*Mean_propotion predicted_* = 0.143) (t(23) = 1.67, *p* = 0 .11). This difference in distance effects between non-symbolic and symbolic formats was statistically significant for both zero (t(23) = 5.45, *p* < .001) and one (t(23) = 3.48, *p* = .002), and is suggestive of more gradedness in the representation of non-symbolic than symbolic numerosities, consistent with previous work describing narrower tuning curves for symbolic numerals^6,13^. We note that a graded RDM still captured a significant portion of variance in the symbolic data (Figure 2C), due to (graded) confusions between non-neighbouring numerosities (e.g., 5 is predicted more often when the classifier sees a 4 than when it sees a 0).

### Representations of Zero are Shared Between Symbols and Empty Sets

Together, our previous analyses establish that neural representations of zero are graded (especially for non-symbolic numerosities) and situated within a number line spanning other countable numerosities from 1 to 5. We next asked whether representations of zero were format- and task-independent – generalising across non-symbolic (empty set) and symbolic (‘0’) stimuli, and across the same/different and averaging tasks. As a first step towards testing for cross-format representations of number, we first computed the Exemplar Discriminability Index^10,41,43^ as a measure of how similar matching cross-format numerosities were (e.g. ‘1’ and one dot, ‘2’ and two dots, etc.) compared to non-matching numerosities (e.g. ‘1’ and five dots). This EDI analysis indicates a significantly higher degree of similarity for matching cross-format numerosities from ∼200ms onwards (Figure 3A).

**Figure 3.**
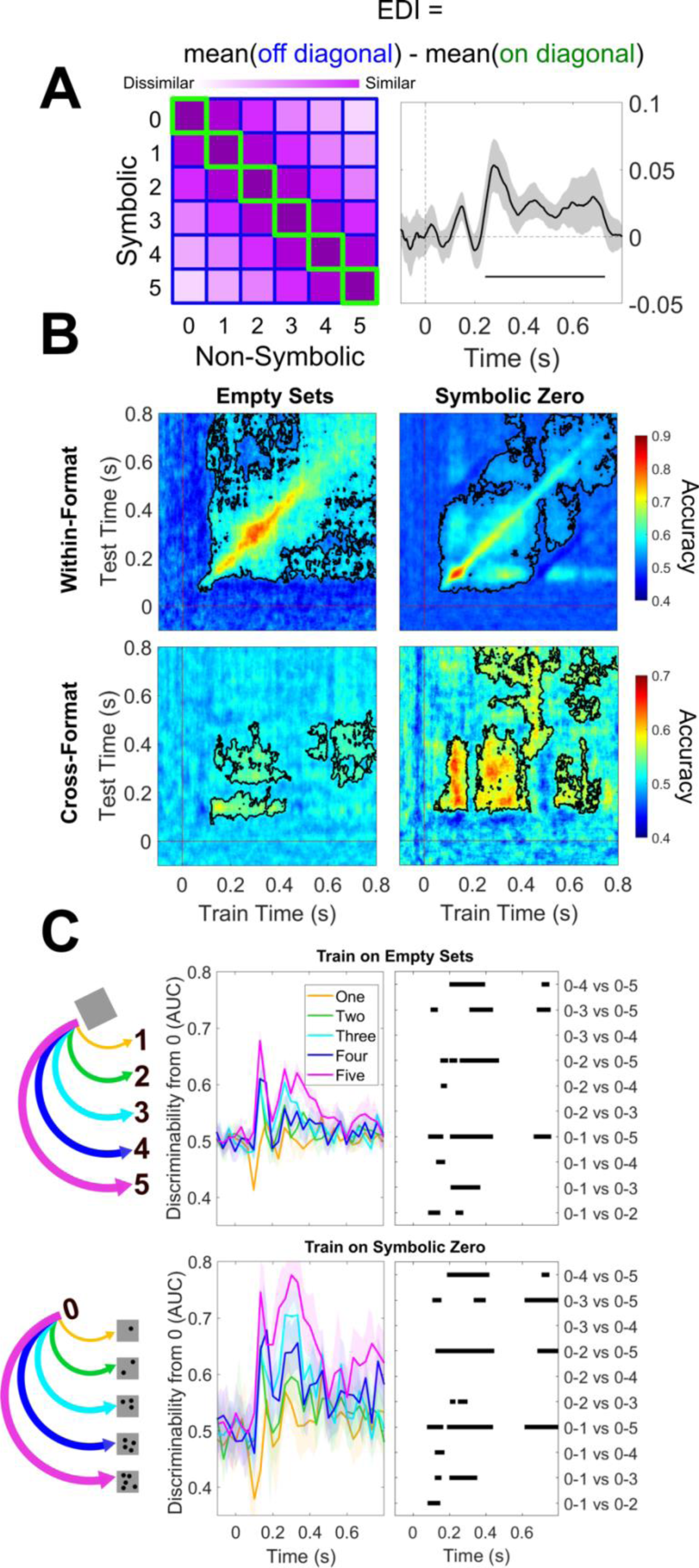
Cross-format, Graded Representations of Numerical Absence. A. The Exemplar Discriminability Index was calculated over time as a measure of the similarity numerosities have to their cross-format counterparts (e.g., testing whether symbolic zero was more similar to empty sets than different dot patterns). Significant EDI was found from around ∼200ms after stimulus onset. B. Representations of numerical absence generalise over numerical format. Top: A decoder trained to decode zero from natural numbers reveals stable representations of zero up to ∼450ms after stimulus presentation for both non-symbolic (left) and symbolic (right) formats, with more dynamic / unstable representations observed towards the end of the epoch. Bottom-left: A decoder trained to decode empty sets also distinguished symbolic zero from non-zero symbolic numerals. Bottom-right: A decoder trained to distinguish symbolic zero from non-zero symbolic numerals also distinguished empty sets from non-symbolic numerosities. Black lines indicate clusters of significantly above chance decoding, p <.05, corrected for multiple comparisons. C. Left: Illustration of the hypothesis that abstract representations of numerical absence are situated on a graded number line that generalises across format, with empty sets represented as more similar to symbolic numeral one than numeral five (top), and symbolic zero as more similar to one dot than five dots (bottom). Centre: Training a classifier to decode non-symbolic empty sets from non-symbolic numerosities and testing it on symbolic numbers in a pairwise manner revealed increasing discriminability as distance from zero increased (top). The same cross-format distance effect is observed when training a classifier on symbolic zero and testing it on non-symbolic numerosities (bottom). Shaded areas represent 95% CIs. Right: clusters of significant differences between different numerosities’ discriminability from zero, p<.05, corrected for multiple comparisons. An increase in discriminability for numbers further from zero reveals a cross-format distance effect.

Next, to specifically test for a cross-format representation of zero, we performed decoding analyses focused on dissociating numerical zero from non-zero numerosities. If a binary classifier trained to distinguish zero from non-zero numerosities in one numerical format is subsequently able to separate zero from non-zero numerosities in another numerical format, this furnishes evidence for an abstract neural representation of numerical absence that is common to both formats.

Decoders trained to distinguish numerical absence within each format separately revealed stable representations of numerical zero from approximately 100ms to 450ms after stimulus presentation, before exhibiting a more dynamic temporal profile until the end of the trial epoch (Figure 3B, top). Crucially, these decoders could also successfully classify representations of zero in the opposing format to which they had been trained (Figure 3B, bottom) – both when generalising from empty sets to the decoding of symbolic numerosities, and when generalising from symbolic zero to non-symbolic dot stimuli. This cross-decoding was successful over the initial 350ms period where the within-format decoders identified stable representations of numerical absence, although generalisation was generally stronger when generalising from symbolic zero to empty sets than vice-versa.

### Graded Representations of Zero are Invariant to Numerical Format

The previous analyses were designed to reveal whether representations of numerical zero generalise across formats, but did not provide a test of whether a binary or graded architecture supports zero representations. To probe the representational structure of cross-format representations of zero, we leveraged the numerical distance effect already identified for within-format representations (Figure 2). To test for such effects, we used a one-vs-one decoding approach to compute the discriminability between zero and each non-zero numerosity in the alternative numerical format (Figure 3C, middle). This approach allowed us to specifically examine how numerical zero is represented with respect to other numerosities. Tests of cross-format representations across all numerosities from zero to five are presented in Supplemental Figure 3 and Supplemental Figure 4. Strikingly, neural representations of symbolic zero (‘0’) were more often confused with one or two dots in the non-symbolic task, than they were with four or five dots (Figure 3C, middle). Similarly, neural representations induced by non-symbolic zero (empty sets) were more often confused with the symbolic numeral 1 or 2 than they were with symbolic numerals 4 or 5. Pairwise tests comparing the discriminability of different non-zero numerosities from zero revealed clusters of significant differences in discriminability (Figure 3C, right), with an increased distance from zero increasing discriminability. Together, these cross-format analyses support a hypothesis that an approximate, graded representation of numerical absence is engaged not only by symbolic zero (‘0’) but also by non-symbolic empty set stimuli.

We also sought to test a more stringent hypothesis that abstract, format-independent neural representations of zero are themselves situated within a cross-format neural number line – thus extending the question of format-independence to now include all numerosities from 0 to 5. A representational dissimilarity matrix situating abstract numerosity representations within a graded number line significantly predicted the neural data (Supplemental Figure 3, left). Testing for cross-format distance effects between all numerosities using RSA also revealed a qualitative distance effect, although this did not reach statistical significance (Supplemental Figure 3, right). Finally, multidimensional scaling of neural representations induced by symbolic and non-symbolic numerosities in a shared space corroborated evidence for a distance effect for zero across tasks (Supplemental Figure 4).

Finally, to explore how similar cross-format representations of zero were to one another, the zero-specific decoders trained in Figure 3B were presented with all the cross-format numerosities from zero to five. The decoder’s decision evidence was taken as a measure of the discriminability of each numerosity from its cross format zero. We found several timepoints where the zero was significantly less discriminable from its cross-format counterpart than any other numerosity (Supplemental Figure 5), suggesting that representations of symbolic zero are most closely related to representations of non-symbolic empty sets than other non-symbolic numerosities, and vice versa.

### Format-Invariant Representations of Numerical Zero are Localised to Posterior Association Cortex

Finally, we sought to localise representations of format-invariant numerical zero in the brain. To do this, we reconstructed and compared source-level neural activity for zero and non-zero numerosities in both the non-symbolic and symbolic tasks. By performing mass-univariate contrasts of broadband source power (zero > non-zero numerosities) in both the non-symbolic (Figure 4A top; peak voxels (*xyz*): left hemisphere = -36, -24, 56; right = 60, -64, -24) and symbolic (Figure 4A bottom; peak voxels (*xyz*): left hemisphere = -28, -56, 32; right = 28, -72, 8) tasks and computing the conjunction between these two contrasts (Figure 4B; peak voxels within conjunction (*xyz*): non-symbolic task: left hemisphere = -20, -64, 32; right = 60, -64, -24; symbolic task: left hemisphere = -28, -56, 32; right = 20, -48, -64), we were able to show that neural activity induced by numerical absence is distributed across the posterior association cortex (Figure 4B). To explore whether the zero representations localised to this region exhibited a graded structure (as found in sensor-level analyses, Figure 3C), we performed multidimensional scaling on source level activity patterns for each numerosity and format. Neural responses to zero within this conjunction map were again situated within a number line populated by non-zero numbers, with numerical magnitude increasing along a single dimension that was similar for both symbolic and non-symbolic formats (Figure 4B, bottom-right).

**Figure 4.**
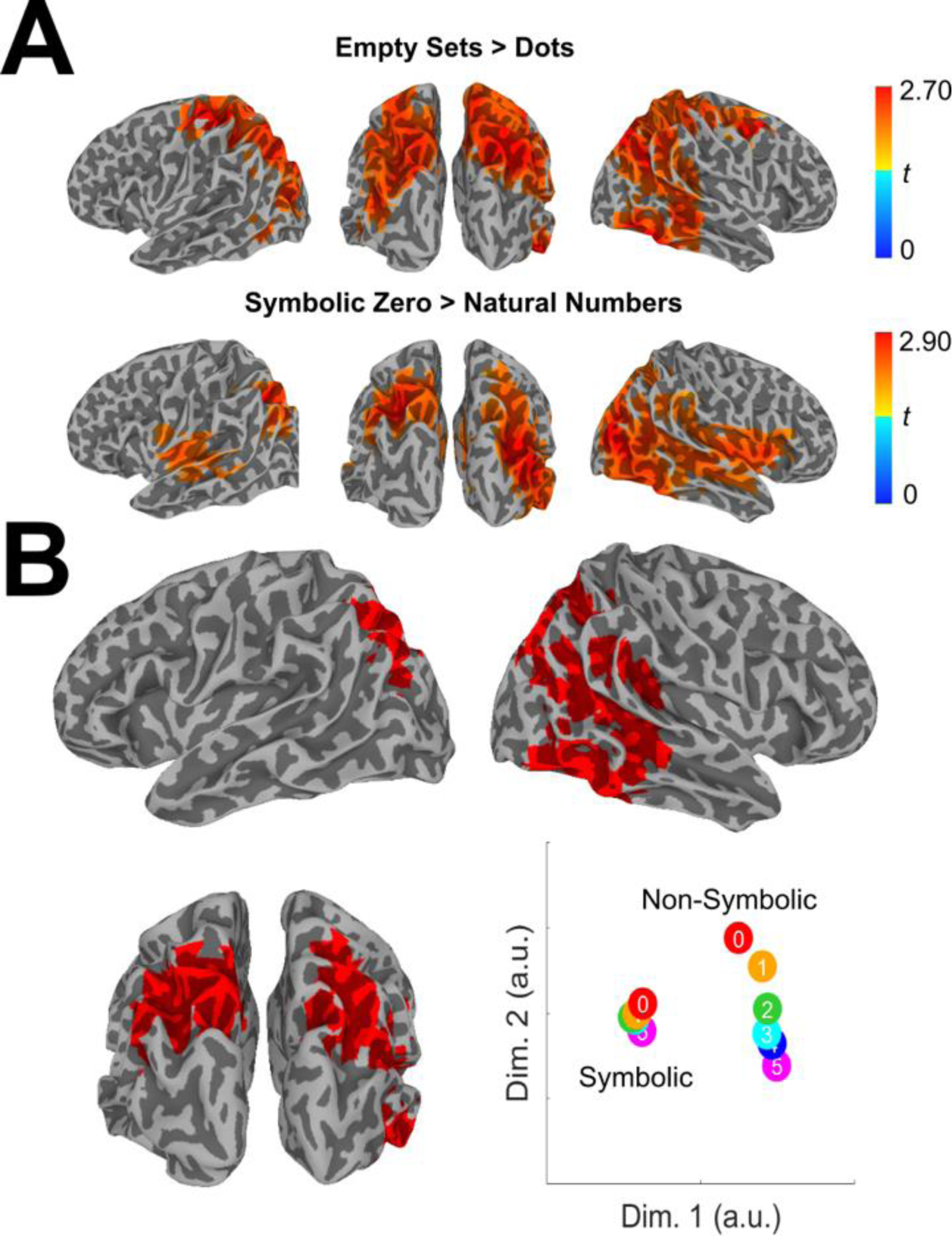
Neural activity induced by numerical zero localised to posterior association cortex. *A*: Mass univariate contrasts of source power revealed regions more active following presentations of zero vs. non-zero numerosities in non-symbolic (top) and symbolic (bottom) tasks. Colour represents t-value and only clusters significant at *p <* .05 are presented, corrected for multiple comparisons. *B*: A conjunction of zero > non-zero contrasts in both numerical formats yielded a map identifying broad regions of the posterior association cortex as representing numerical absence across numerical formats. Multidimensional scaling of each numerosity’s neural pattern within these regions revealed a graded representational structure of numerical magnitude along a single dimension that was similar for both formats.

## Discussion

The number zero is associated with unique psychological properties compared to natural numbers. Here, we characterise the neural representation of numerical zero in the human brain. We describe how numerical zero occupies a slot at the lower end of neural number lines for both symbolic and non-symbolic numerical formats. We go on to show that a component of this representation is both task- and format-independent, such that empty sets – the absence of dots – generalised to predict the neural profiles and distance effects observed for symbolic zero. These abstract, format-invariant representations of zero were situated at the lower end of a neural code for number that was localised across the posterior association cortex.

Our finding that representations of numerical absence have a format-invariant component extends previous work documenting neural representations of numerosity that generalise across countable non-symbolic elements and their symbolic counterparts ^13,14,33^. Here we show how neural representations of non-symbolic empty sets, which do not contain any countable items, also share variance with symbolic zero (Figure 3) across qualitatively different tasks with distinct behavioural profiles. These abstract representations of zero were localised to regions of the posterior association cortex that have previously been associated with numerical processing^12,14,18,35^.

That zero is situated at the lower end of a neural number line in the human brain is consistent with an emerging body of work examining representations of zero in non-human animals^22–24^. Across two different studies, single neurons selective for non-symbolic empty sets were found in the parietal and prefrontal cortex of non-human primates^23,24^. In line with the present results, many of these neurons – but not all – were found to exhibit distance effects with non-zero numbers. When comparing non-symbolic and symbolic instances of zero, we found symbolic instances were more discrete and less graded than non-symbolic instances, consistent with work describing sharper tuning curves for symbolic number representations^6,13^. Recent single-cell recordings in the human medial temporal lobe have also identified discrete symbolic zero-selective neurons that did not exhibit graded activations in relation to non-zero symbolic numerals^29^. Strikingly, however, the majority of our analyses revealed a graded representation of zero that generalised across both symbolic and non-symbolic formats. This is in keeping with 7T fMRI data showing that neural populations at the lower boundary of numerically-tuned topographic maps exhibit a monotonic decrease in response to increasing numerical magnitude^31^, a finding suggestive of evidence for neural populations tuned to numerosities below one.

We took care to ensure that the neural representations of zero identified in our data were not trivial consequences of zero being classified as the ‘lowest’ stimulus in our tasks. The concern here is that if our tasks required participants to adopt a particular mathematical attitude towards zero, then decoding of this task-dependent concept would confound any results aimed at identifying task-invariant representations of numerical absence. We consider this explanation of our results as unlikely, however, as, by design, the symbolic and non-symbolic tasks required adopting qualitatively distinct mathematical attitudes towards zero stimuli: the match-to-sample task necessitated deciding whether two dot stimuli were the same or different, whereas the symbolic task required maintenance of condition-specific numerical averages. Because the non-symbolic task did not require participants to order stimuli, any format-invariant representations of zero cannot be explained by a generic requirement to identify lower vs. higher numerosities. Despite these task differences, explicit numerical processing of the stimuli was common to both tasks, and by design, both of these tasks afford the locking of the MEG data to the onset of examples of zero (and other numerical) stimuli that are embedded in a wider task context.

In both cross-decoding (Figure 3; Supplemental Figure 5) and RSA (Supplemental Figure 3) analyses, we observed cross-format representations emerging initially between 100-200ms post-stimulus before diminishing and re-emerging approximately 300ms after stimulus presentation. The small number of previous studies which have explored the time course of abstract numerical representations in the human brain document generalisation from symbolic formats to non-symbolic formats or alternative magnitude domains from ∼300ms onwards^10,15^, consistent with this later peak in our data. The earlier peak in contrast is consistent with other studies documenting decodability of non-numerical conceptual representations (such as animate/inanimate and artificial/natural) as early as ∼100ms post-stimulus^58,60^. It remains debated whether findings of format-invariant numerical codes are explained by single neurons coding for the same numerosities across formats, or whether they reflect the recruitment of neighbouring format-specific neural populations that are interdigitated within a numerosity map^36^. Future intracranial recording studies will be required to determine whether single cells in the human brain code for numerical zero in both non-symbolic and symbolic formats. However, our finding of cross-format distance effects is more consistent with a shared neural code, as it is less likely that spatially overlapping but format-specific neural codes would also generalise to exhibit cross-format distance effects with more distant numerosities. Furthermore, our finding of a cross-format code for numerical zero is in keeping with behavioural studies that have found format-invariant distance effects for both symbolic zero and empty sets^53^.

Behavioural tasks have previously been used to investigate zero in relation to a mental number line^19,21,47,48^. For example, the SNARC (spatial-numerical association of response codes) effect extends to numerosity zero, with faster response times when zero responses were given with the left hand^19^. Similarly, the size-congruity effect^49^ has been exploited to suggest that zero occupies a definite position on a mental number line: responses to zero are facilitated when it is physically smaller than an alternative numeral^21^. Notably, ‘end effects’ have been established in size congruity paradigms, whereby stimuli perceived to be at the ‘end’ of the number line exhibit facilitated response times. End effects have been found for symbolic zero even when presented amongst negative numbers^21^ and in response to non-symbolic empty sets during numerical comparison tasks^20^, suggesting both symbolic and non-symbolic zero may be situated at the beginning of a mental number line. Such findings obtained using elegant behavioural assays are consistent with our findings that numerical zero is incorporated into a graded neural number line amongst other non-zero numbers (at least in some contexts^51,53^).

Classical accounts of how symbolic representations of number are mapped to their non-symbolic counterparts do not readily explain how the symbolic concept of zero is generated from non-symbolic empty sets. These models describe a neural architecture in which specific numerosities are mapped onto numerosity-tuned neural populations that form a neural number line, or place-coding system^14,65^. However, according to such models, non-symbolic stimuli only proceed to this place-coding system via a summation procedure, where an increase in the number of non-symbolic elements accumulate a greater degree of activation^66–69^. A summation explanation therefore fails to account for how non-symbolic empty sets may be mapped to symbolic zero, since empty sets offer no countable elements to accumulate^21^. Computational models of object recognition have, however, documented the emergence of spontaneous representations of non-symbolic zero without any training on numerical stimuli^70^, suggesting that the concept of zero can be readily acquired from statistical regularities in visual input without necessarily relying on a summation procedure.

A shared neural representation underpinning both non-symbolic empty sets and symbolic zero is consistent with recent suggestions that representations of numerosity zero may have emerged from more fundamental representations of sensory absence^2^. On this account, low-level perceptual representations indicating an absence of sensory stimulation^62,63^ provide the perceptual grounding for a conceptual representations of numerical zero^2^ – consistent with a broader principle that the human brain co-opts basic sensory and motor functions in the service of more complex cognitive abilities^38^. Such a hypothesis aligns with similar behavioural signatures for the processing of absence across perceptual and numerical domains. For instance, reading times are increased for number zero compared to non-zero numbers^39^, whilst reaction times for deciding a stimulus is absent are higher than for deciding a stimulus is present^40,42^. Additionally, judgements about the absence of features mature later in children than judgements about presence^44,45^, a developmental pattern mirrored by the late mastery of numerical zero in children ^5,46,47^. We note however that the neural responses recorded in our study to empty-set stimuli were still within the context of a numerical task – and, as such, our results are specific to the concept of numerical absence and do not provide a direct test of a link between numerical zero and sensory absence. Out study offers a path towards a formal test of this hypothesis in future work - for instance, investigating relationships between numerical absence and non-numerical perceptual absences^40,62,64^.

The adoption of the number zero has enabled great advances in science and mathematics ^1^. Here, we show that the human brain represents this unique number by incorporating representations of numerical absence into a broader neural coding scheme that also supports countable numerosities. Representations of numerical zero were found to be format-invariant and graded with respect to non-zero numerosities, and were localised to regions of the posterior association cortex previously implicated in numerical cognition. Our results demonstrate that neural number lines include zero, and, more importantly, provide initial evidence that the abstract concept of symbolic zero is linked to representations of non-symbolic empty sets. Our study lays the foundations for a deeper understanding of how the human ability to represent the number zero may be grounded in perceptual capacities for detecting an absence of sensory stimulation.

## Acknowledgements

We would like to thank the Imaging Support Team at the Wellcome Centre for Human Neuroimaging for their help with data collection. B.B is supported by the Leverhulme Trust Doctoral Training Programme for the Ecological Study of the Brain [DS-2017-026]. S.M.F. is a CIFAR Fellow in the Brain, Mind and Consciousness Program, and supported by a Wellcome/Royal Society Sir Henry Dale Fellowship [206648/Z/17/Z] and a Philip Leverhulme Prize from the Leverhulme Trust. The Wellcome Centre for Human Neuroimaging is supported by core funding from the Wellcome Trust [206648/Z/17/Z].

## Author Contributions

B.B. conceived and performed experiments, carried out analyses, visualised and interpreted results, wrote the first draft of the manuscript, and edited later drafts. S.M.F conceived experiments, provided resources and supervision, secured funding, and reviewed and edited the manuscript.

## Declaration of Interests

The authors declare no competing interests.

## Supplemental Information

Document S1: Supplemental Figures 1 to 5.

## Methods

### Data and code availability

All original MEG data has been deposited at OSF (https://osf.io/vr7qp/) and is publicly available. DOIs are listed in the key resources table.

All original code has also been deposited at OSF (https://osf.io/vr7qp/) and is publicly available. DOIs are listed in the key resources table.

Any additional information required to reanalyse the data reported in this paper is available from the lead contact upon request.

### Study Participant Details

Twenty-nine participants (*M*_age_: 29.27 years, *SD*_age_: 10.69) took part in the MEG experiment at the Wellcome Centre for Human Neuroimaging, University College London. Five participants either failed to follow task instructions (chance performance on one or more tasks) or did not complete the experiment and were therefore excluded from analysis. All analysis was performed on the remaining sample of 24 participants. Informed consent was given before the experiment and ethical approval was granted by the Research Ethics Committee of University College London (#1825/005).

### Method Details

#### Stimuli

Numerical dot stimuli were created using custom MATLAB 2021b (Mathworks) scripts and consisted of different numbers of dots (from zero to five) on grey backgrounds (Figure 1C). There were two sets of dot stimuli, a standard set and a control set. In the standard set, dot size was pseudorandomly specified, and each dot was pseudorandomly located around the centre of the screen. In the control set, low level visual properties of the stimuli (total dot area, density, luminance) were constant across numerosities. Total dot area was controlled for by systematically reducing the size of the dots as the number of dots increased, such that the total number of pixels included in a stimulus were constant across numerosities. To control for density, dots were located within an invisible circle, with the radius of the circle determined by the number of dots. Larger numerosities had larger circles, thereby ensuring that patterns with more dots were not systematically denser. Finally, in both stimulus sets, 50% of dots were black and 50% were white, such that the contrast of the dot patterns did not increase with numerosity (the increase in contrast generated by an increasing number of black dots was cancelled out by the increased number of white dots). Supplementary Figure 1 reports the correlation between these non-numerical features and numerosity in the controlled stimulus set, highlighting how our stimulus-generating procedure successfully controlled for the association between low-level visual properties and numerosity. Empty set stimuli contained only a grey background in both stimulus sets.

To help prevent participants relying on low level visual cues in identifying empty set stimuli, the background luminance was varied within and across stimulus sets and the background square size was randomly varied across all stimuli. The two stimulus sets allowed us to test whether numerical representations generalised across controlled and uncontrolled stimulus sets, which if successful indicates that numerosity representations are not merely picking up on low-level stimulus features correlated with number. Indeed, a control analysis confirmed that numerical information was extracted from the stimuli independently from physical features (see Representational Similarity Analysis; Supplemental Figure 3). As such, both standard and control stimulus sets were included in the analysis of the non-symbolic data.

#### Experimental Procedure

The tasks were presented to subjects using MATLAB (Mathworks) and the Psychophysics Toolbox^71,72^. Participants practiced the tasks on a computer before the MEG session. In the MEG scanner, the tasks were performed in alternating miniblocks with 35 symbolic trials and 54 non-symbolic trials per MEG recording block. The order of the tasks would swap on each block, and the starting order was counterbalanced across participants. There were 9 MEG blocks in total, resulting in 315 symbolic numeral trials and 486 non-symbolic dot trials across the whole experiment. Participants responded using two buttons on a button box and their right thumb.

##### Non-symbolic Task

Participants performed a match to sample task on dot stimuli^22,24^. On each trial, participants saw a sample image containing between zero and five dots for 250ms followed by a fixation cross for 800ms. A test image, also containing between zero and five dots, was then presented for 250ms, followed before another 800ms fixation period (Figure 1A). Within a trial, a single stimulus set was used for both the sample and test image. Participants reported whether the number of dots in the test stimulus matched that of the sample stimulus, or not. The response was followed by feedback in the form of a coloured rectangle surrounding the response options, with green and red used to indicate correct and incorrect answers, respectively. Response options were positioned randomly on each trial to eliminate any correlation between the decision and motor response. Intertrial intervals were also sampled randomly from a uniform distribution between 500-1000ms.

##### Symbolic Task

We adapted the symbolic numeral averaging task introduced by^10^ to include the number zero. In one trial, ten numerals ranging from zero to five were presented in a random order (Figure 1B). Five of the numerals were blue and five were orange. Each numeral was displayed for 250ms with an interstimulus interval of 100ms. The numerals were randomly selected on each trial to obey the constraint that the mean of the blue numerals could not equal the mean of the orange numerals. The response required at the end of each trial was counterbalanced across subjects, with half of the subjects reporting which set of numerals (orange or blue) had the highest average, and the other half reporting the set with the lowest average. Participants had 2000ms to respond, after which they were given feedback in the form of a green (correct) or red (incorrect) rectangle surrounding the response options. Again, to disentangle participants’ decisions from motor responses, response options were positioned randomly on each trial. Intertrial intervals were randomly sampled from a uniform distribution between 500-1000ms.

#### MEG Preprocessing

MEG data were analysed using FieldTrip^50^. MEG was recorded continuously at 600Hz using a 273-channel axial gradiometer system (CTF Omega, VSM MedTech) while participants sat upright inside the scanner. To remove line noise, the raw MEG data were preprocessed with a Discrete Fourier Transform and bandstop filter at 50Hz and its harmonics. The numeral task was segmented into epochs of -500ms to 4000ms relative to trial onset. For the dot task, the segments were from -200ms to 2500ms. Baseline correction was performed where, for each trial, activity in the pre-trial window was averaged and subtracted from the entire epoch per channel. The data were downsampled to 300Hz to conserve processing time and improve signal to noise ratio. During artefact rejection, trials with high kurtosis were visually inspected and removed if they were judged to contain excessive artefacts. To assist in removing eye-movement artefacts, an independent components analysis was carried out on the MEG data, and the components with the highest correlation with eye-tracking data were discarded after visual inspection. Components showing topographic and temporal signatures typically associated with cardiac artefacts were also removed by eye. This procedure was performed separately for the numeral and dot task. Finally, a second stage of epoching was performed to generate trials of individual numerosities. In the numeral task, trials were segmented into -100ms to 800ms epochs around each numeral onset. Trials were then baseline corrected again using the pre-stimulus window. In the dot task, trials were segmented into two different -200ms to 800ms epochs with respect to the onsets of the sample and test stimuli. All analyses used the sample images only. Finally, all analyses focusing on shared representations across notational formats were performed on the shared timepoints of -100ms to 800ms relative to stimulus presentation.

### Quantification and Statistical Analysis

#### Representational Similarity Analysis

Representational Similarity Analysis (RSA) allows us to test specific hypothesis about how neural representations are structured^52^. Here, we tested for the existence of a distance effect across numerosities (Figure 2C). To do this, we defined a model representational dissimilarity matrix (RDM) that describes the dissimilarity of two numerosities as a function of their numerical distance. To compare this model dissimilarity matrix with the neural data we first created a neural dissimilarity matrix that represents the similarity in neural patterns associated with each numerosity. To do this, we first ran a linear regression on the MEG data with dummy coded predictors for each of the six numerosities (trial numerosity coded with a 1, alternative numerosities coded with a 0). This produced a coefficient weight for each numerosity at each time point and sensor. These weights were then combined into a vector, representing the multivariate neural response for each numerosity, averaged over trials. To create the neural RDM, we computed the Pearson distance between each pair of condition weights over sensors, resulting in a 6×6 neural RDM reflecting the pairwise similarity of neural patterns associated with each numerosity. These neural RDMs were smoothed over time via convolution with a 60ms uniform kernel. To compare the neural and model RDMs, at every time point we correlated the lower triangle of each matrix (excluding the diagonal) using Kendall’s Tau rank correlation^54^.

Cross-task RSA was performed in the same manner, except here there were 12 predictors in the linear regression (0-5 symbolic, 0-5 non-symbolic). This resulted in a 12×12 neural RDM, of which we used the quadrant representing the cross-task pairwise similarities between numerosities when comparing with the model RDM (Supplemental Figure 3). The whole quadrant including the diagonal was used in this analysis. This is because here the diagonal does not contain redundant information, but rather the similarity of the same numerosity across two different formats, and cells in the upper triangle represent different pairwise similarities to those in the lower triangle.

Finally, to test whether numerical information was decodable from non-symbolic stimuli over and above the physical features of the stimuli, we ran a cross-stimulus set RSA in the same manner as above, except now we tested exclusively within the non-symbolic task (Supplemental Figure 2). As such, the 12 predictors were: 0-5 from the standard set and 0-5 from the control set. This RSA established whether representations of numerical magnitude generalised across stimulus set, and therefore went beyond information that could be derived solely from physical features of the stimuli.

#### Decoding Analyses

To examine the representational structure of the number zero more specifically across symbolic and non-symbolic formats, we employed different decoding techniques using both multiclass and binary decoders. First, to reveal the temporal profile of numerosity representations, we trained a multiclass Linear Discriminant Analysis (LDA) decoder to decode numerosities zero to five (Figure 2B). This was performed in a temporal generalisation procedure, whereby the classifier was trained on each time point and tested on all other time points^27^. This process results in a train time x test time decoding accuracy matrix, which illustrates how stable representations of numerosity are over time.

We performed both within-format and cross-format decoding procedures. Within-format decoding involved training and testing a classifier to identify numerosities on trials from one format (e.g. numerals or dots). In cross-format decoding, we trained the classifier on one format and tested it on the other (e.g., training on symbolic trials and testing on non-symbolic trials, and vice versa). For the within-format approach, we implemented a 5-fold cross-validation strategy. Prior to decoding, five trials per numerosity were averaged and the resulting average trials was balanced per numerosity. It is worth noting that cross-validation is not required in cross-format decoding because the test data is never seen by the classifier during training, and thus there is no risk of overfitting. Cross-format decoding allows us to empirically assess whether the neural patterns associated with numerals share a common neural code across formats.

To complement our RSA analyses and isolate the representational structure underpinning numerical zero specifically, we extracted the confusion matrices from the decoders (Figure 2A, 2D). Confusion matrices indicate how often different stimulus classes (i.e., numerosities) are confused for one another, and this information can be used to infer the organisation of neural representations. For example, a decoder that confuses zero with the number one more than the number two displays evidence for a numerical distance effect. The data used to train the decoders from which these confusion matrices were extracted was time-averaged over the timepoints where the initial multiclass decoder could decode numerosity significantly above chance (non-symbolic: 70ms – 800ms, symbolic 56.7ms – 800ms; Figure 2B). We also computed confusion matrices across time (Figure 2A).

To examine whether representations of zero could reliably be dissociated from numerosities presented in the alternative format, we created a decoding procedure using a binary LDA classifier to decode zero vs. non-zero numerosities (Figure 3B). Within this training regime, the number of trials per non-zero numerosities was kept equal, and the number of zero trials vs. non-zero numerosity trials was also balanced. The resulting ‘zero’ decoder was uniquely trained to identify neural representations of numerical zero in symbolic or non-symbolic format and was tested on the other format to identify format-invariant representations of zero.

Finally, to reveal whether abstract representations of numerical zero exist on a graded number line, or whether they are categorically distinct from other numbers, we ran a new cross-format decoding analysis using binary classifiers. Here, we trained the decoders to discriminate zero vs. all non-zero numerosities (one to five) separately, and then tested these binary decoders on the corresponding numerosities in the opposite format. This resulted in five different classifiers per format. Specifically, we trained five different decoders to dissociate: symbolic zero vs symbolic one, symbolic zero vs symbolic two, symbolic zero vs symbolic three, symbolic zero vs symbolic four, and symbolic zero vs symbolic five. We then tested these decoders on empty sets vs one dot, empty sets vs two dots, empty sets vs three dots, empty sets vs four dots, and empty sets vs five dots, respectively. This was also done in the reverse direction: training on non-symbolic trials and testing on symbolic numerals. We used the area under the receiver operating characteristic (AUROC) as a metric for discriminability between each pair of classes. In line with the hypothesis that format-invariant representations of zero exist on a graded, abstract neural number line, we expected the discriminability to improve as the numerical distance from zero increased (Figure 3C). To statistically test whether this was the case, we performed one-tailed, paired comparisons between the discriminability of successive numbers with zero (e.g., by comparing 0-2 vs. 0-1, 0-3 vs. 0-2, etc.; Figure 3C).

For all decoding analyses, we utilized multiclass or binary LDA decoders in conjunction with the MVPA-light toolbox^55^ integrated with FieldTrip. To improve the robustness of the classifier, we applied L1-regularization to the covariance matrix, and the shrinkage parameter was automatically determined using the Ledoit-Wolf formula within each training fold^73^.

#### Source Reconstruction

Both FieldTrip’s template single shell head model and its standard volumetric grid (8mm resolution) were warped to participants’ individual fiducial points, generating a subject-specific forward model aligned in MNI space. Source reconstruction was performed using a linearly constrained minimum variance (lcmv) beamformer^56^ which applies spatial filters to the MEG data to generate source-level time courses. To reduce the impact of noise on the source estimates, we used a regularisation parameter of lambda = 5%. For each task, spatial filters were calculated by combining the leadfield matrix with the data covariance matrix across all numerosities and the timepoints coinciding with the stable cluster of significantly above-chance decoding in the zero vs. non-zero cross-task classifier (100 – 450ms). These spatial filters were then applied to zero trials and non-zero trials separately, generating reconstructed maps of source activity for these two trial types. We contrasted the broadband source power of zero > non-zero trials in a mass-univariate procedure across subjects for each task separately (Figure 4A) with an alpha parameter of p < .05, corrected for multiple comparisons. For binary LDA classifiers, this is equivalent to localising the classifier weights^57^, and therefore gives an indication of which brain regions drove our decoding results. We computed the conjunction of these two contrasts, revealing the voxels where zero stimuli were dissociable from other numbers in both symbolic and non-symbolic formats (Figure 4B).

Multidimensional scaling of source space activity was performed using the same beamforming parameters to calculate spatial filters over combined non-symbolic and symbolic trials. Using these filters, virtual channels were created for each source location within the map defined by the conjunction analysis. The virtual channels were then used to create a cross-task representational dissimilarity matrix in the same manner as described for the cross-task RSA sensor-level analysis. This was then submitted to MATLAB’s cmdscale function for multidimensional scaling.

#### Statistical Inference

Across sensor and source level analyses, cluster-based permutation testing was used to statistically test hypotheses and correct for multiple comparisons^59^. For all analyses (decoding, RSA, and source-level contrasts), 1000 permutations were used with cluster-forming alpha parameter of .05 and a significance threshold of .05. It is important to emphasize that this cluster-based permutation testing approach does not provide information about when neural representations emerge. This limitation arises because the statistical inference process does not focus on individual time points; instead, it relies on cluster-level statistics that encompass multiple time points^61^.

## Notes

### Competing Interest Statement

The authors have declared no competing interest.

### Summary of Updates

Discussion updated. Figure 3 updated. Supplemental Analyses added.

